# Comparative Study between In-silico and Clinical Works on the Control of Blood Glucose Level in People with Type 1 Diabetes using Improved Hovorka Equations

**DOI:** 10.1101/2022.09.23.509189

**Authors:** Nur Amanina Mohd Sohadi, Ayub Md Som, Noor Shafina Mohd Nor, Sherif Abdulbari Ali, Mohd Aizad Ahmad

## Abstract

**Background:** Hovorka model is one of the diabetic models which is widely used in the artificial pancreas device (APD) also known as closed loop system, meant for people with type 1 diabetes (T1D). Previous workers had modified some equations in the sub-sections of the Hovorka model, which is also known as improved Hovorka equations, in regulating the blood glucose level (BGL) within normoglycemic range (4.0 to 7.0 mmol/L). However, the improved Hovorka equations have not been tested yet in terms of its usability to regulate and control the BGL in safe range for two or more people with T1D. This study aims to simulate their BGL with meal disturbances for 24 hours using the improved Hovorka equations.

**Methods:** Data for people with T1D were obtained from Clinic 1, Clinical Training Centre (CTC), UiTM Medical Specialist Centre, Sungai Buloh, Selangor. Data collected include gender, age, body weight, mealtimes, meal amount, and duration. Three patients whose ages range from 11 to 14 years old were selected. All patients consumed three meals daily: breakfast, lunch, and dinner. The simulation (in-silico work) was done using MATLAB software, and the BGL profile from both in-silico and clinical works were compared and analysed.

**Results:** It was revealed that the BGLs for all three people with T1D were far better in the in-silico work compared to the clinical work. The BGL for patient 1 was able to achieve normoglycaemia 73% of the time in the in-silico work. Meanwhile, patient 2 managed to stay in the normoglycemic range for 85% of the time in the in-silico work compared to clinical work, which was merely 31%. For Patient 3, the time duration spent in the normoglycemic range was only 16% in the in-silico work compared to none as in the clinical work. The p-values obtained in the study were less than 0.05, indicating that the in-silico work using the improved Hovorka equations was acceptable for predicting the BGL for people with T1D.

**Conclusions:** It can be concluded that the improved Hovorka equations are reliable in simulating the meal disturbances effect on BGL and increasing people in T1D times’ duration in the normoglycemic range compared to the clinical work.

## Introduction

Diabetes is a chronic disease that is characterised by high blood glucose level (BGL). It occurs when beta-cells in the pancreas produce insufficient insulin or cells are unable to utilise insulin effectively. Insulin is crucial in blood glucose regulations by promoting glucose uptake, consequently lowering the BGL [1]. BGL should be regulated within the normoglycemic range between 4.0 to 7.0 mmol/L [2]. Maintaining BGL in a safe range can help prevent long-term complications related to hyperglycaemia (high BGL) and hypoglycaemia (low BGL). A lack of insulin production causes T1D thus disrupts blood glucose regulation. T1D commonly occurs in children and adolescents. The incidence of T1D is more prevalent in Northern Europe than in the rest of the world. The International Diabetes Federation reported 977 cases of T1D in Malaysian children aged 0 to 19 years old in 2019 [3]. People with T1D must monitor their BGL via finger pricks and take either self-monitoring of blood glucose (SMBG) or multiple daily injections (MDI) of insulin to regulate their BGL.

Current practice in blood glucose management is invasive, which causes distress to patients. Thus, the introduction of an artificial pancreas device (APD), also known as a closed loop system, has improved the quality of life of people with T1D for the last few decades. Despite the recent advancement of APD, the control algorithms used are however still lagging in delivering the proper insulin dosage to people with T1D. Previous workers had modified some equations in the sub-sections of the Hovorka model [4], which is also known as improved Hovorka equations, in regulating the blood glucose level (BGL) within normoglycemic range (4.0 to 7.0 mmol/L)[5-6]. However, the improved Hovorka equations have not been tested yet in terms of its usability to regulate and control the BGL in safe range for two or more people with T1D. This study aims to simulate their BGL with meal disturbances for 24 hours using the improved Hovorka equations. Then, the patients’ BGL from the simulation (in-silico work) are compared with the clinical work. This study is a continuation of work from the previous studies [7-9].

## Methods

### Data Collection for Clinical Works

Ethical approval for this study was granted by the UiTM Ethics Committee before data collection was commenced. Information sheets were provided, and formal consent for participation was obtained from parents or guidance of the focus groups, i.e., three people with T1D ages range from 11 to 14 years old. Data collected include name, age, gender, body weight, meal times, amounts of meals consumed, and meal duration. Tables 1 to 3 summarise the amounts of meals consumed in 24 hours for the three people with T1D. Detailed meal contents for the study were adopted from the previous work [10]. All the three people with T1D were on self-monitoring of blood glucose (SMBG) via finger pricks and their BGLs were taken before and after each meal intake namely: during breakfast, lunch, and dinner.

### A Mathematical Model for In-silico Work

The improved Hovorka model based on Hovorka model [4] is specifically designed for people with T1D. The model has two inputs: meal disturbances and bolus insulin. It consists of three subsystems: the glucose subsystem, insulin subsystem, and insulin action subsystem.

Equations for glucose subsystem are as shown in Eq. (1) and (2):

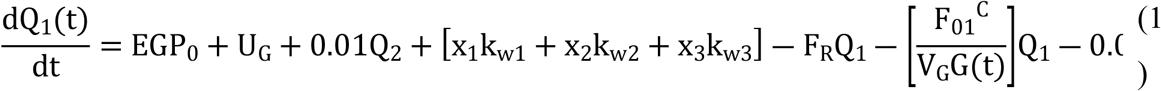

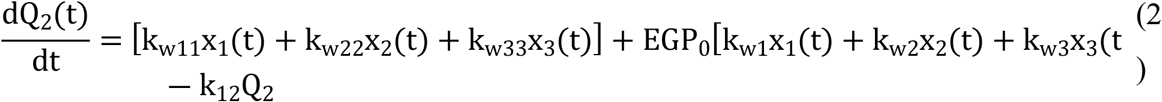

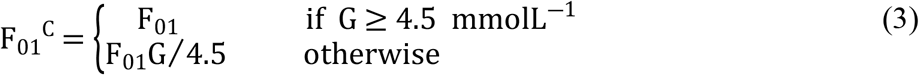

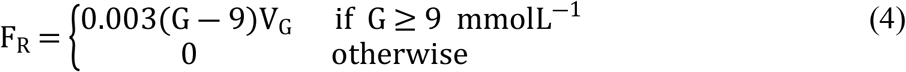

where: Q_1_ (mmol) is mass of glucose in accessible compartment, Q_2_ (mmol) is mass of glucose in non-accessible compartment, k_w1_, k_w11,_ k_w2_, k_w22_, k_w3_, and k_w33_ (min^-1^) are activation rates, k_12_ (min^-1^) is transfer rate, EGP_0_ is endogenous glucose production (EGP) extrapolated to zero insulin concentration, V_G_ (L/kg) is glucose distribution volume, G (mmol/L) is glucose concentration, x_1_, x_2_, and x_3_ are the effects of insulin on glucose transport and distribution, glucose disposal, and EGP respectively, F_01_ ^c^ (mmol min^-1^ kg^-1^) is total non-insulin dependent glucose flux, and F_R_ (mmol/min) is renal glucose clearance.

When a meal is consumed, the carbohydrate (CHO) content will be broken down into glucose prior to being converted into energy. Thus, increasing the BGL. The equation to observe the effect of meal disturbances on BGL is shown in Eq. (5).

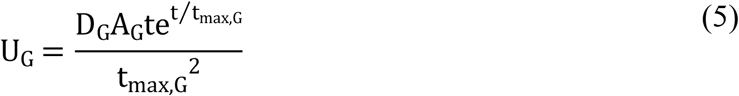

Where,

D_G_ = amount of carbohydrate (CHO) digested (mmol)

A_G_ = carbohydrate bioavailability

t_max,G_ = time-to-maximum of CHO absorption (min)

Eq. (6) to (8) show the insulin subsystem. Insulin is administered subcutaneously. This method is less invasive than the intravenous method, in which insulin is injected directly into the bloodstream.

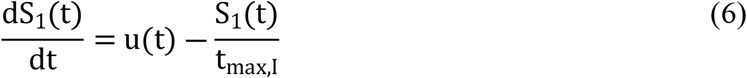

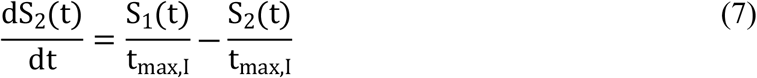

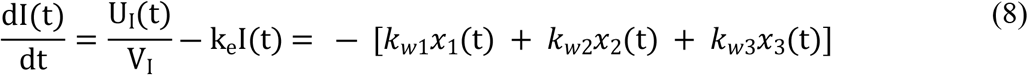

The insulin absorption rate in the bloodstream is given by Eq. (9).

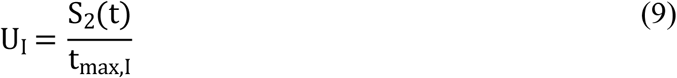

where: S_1_ and S_2_ (mU) are insulin sensitivity in accessible and non-accessible compartments respectively, u(t) (mU/min) is bolus insulin, t_max,I_ (min) is time-to-maximum of insulin absorption, U_I_ (mU/min) is insulin absorption rate, V_I_ (L/kg) is insulin distribution volume, and k_e_ (min^-1^) is fractional elimination rate.

The insulin action subsystem is represented as in Eq. (10) to (12). The insulin-glucose interaction can be observed in this subsystem.

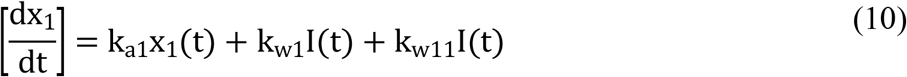

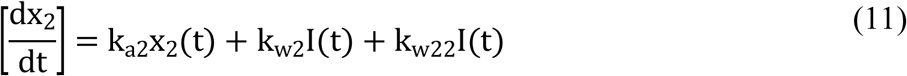

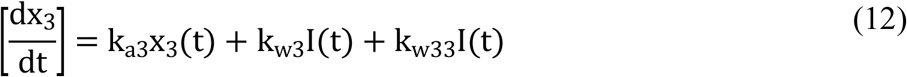

Constants and parameters involved in the equations are shown in the Tables 4 and 5. A diagram of the Hovorka model with improved equations is illustrated in Figure 1.

**Figure 1.**
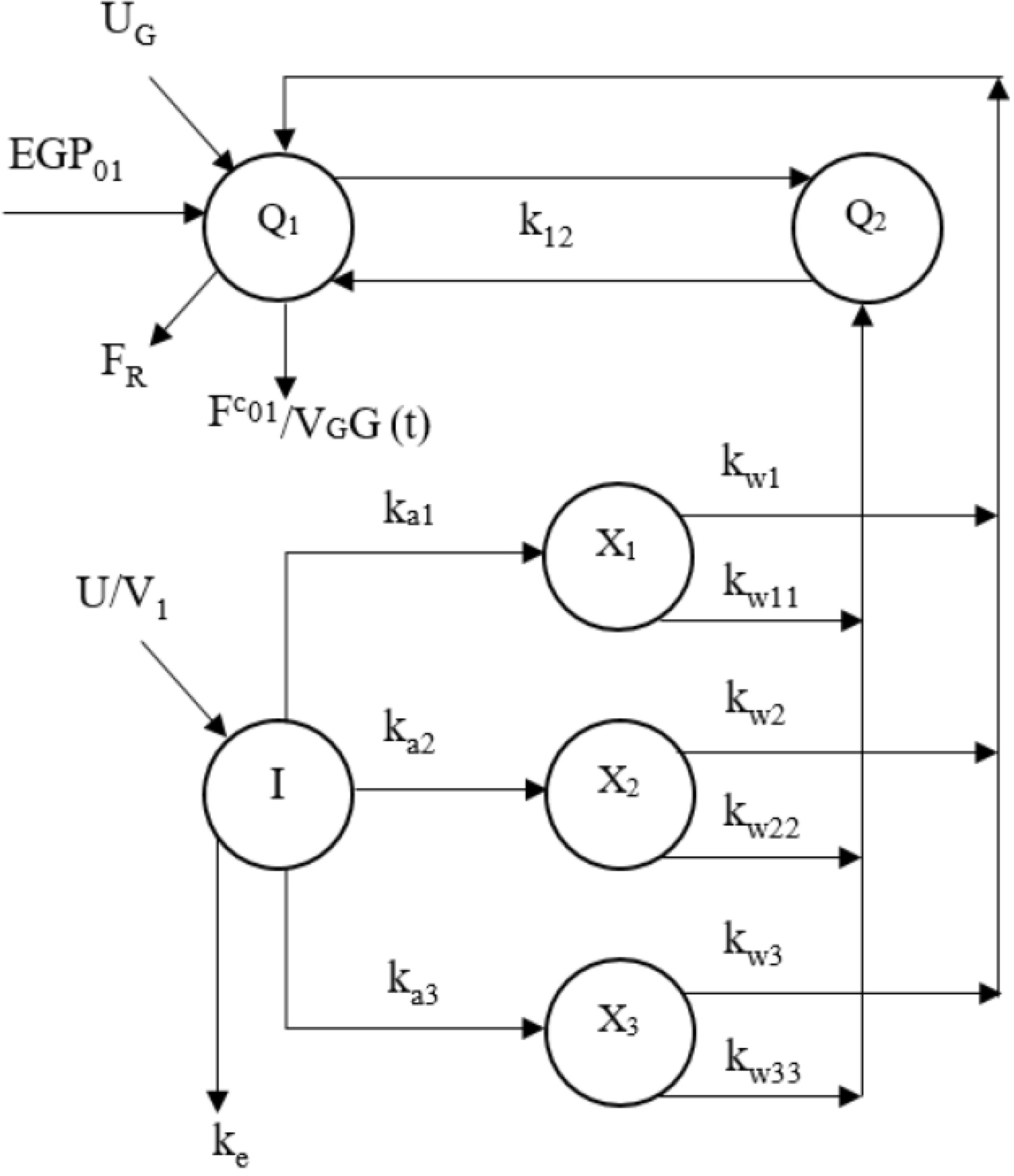
Improved Hovorka equations

The in-silico work was performed based on the estimated meal disturbances on the people with T1D as adopted from [10]. The amounts of insulin administered (U_t_) selected for this study were namely: 0.100 U/min for high dosages, 0.050 U/min for medium dosage and 0.0167 U/min for low dosage as also adopted from [10]. The simulation (in-silico work) using the improved Hovorka equations was done in MATLAB and data obtained were then collected for further analysis and evaluation.

### Sample calculation of CHO intake for patient 1

In addition, sample calculations for patient 1 with a body weight of 27 kg are as follows:

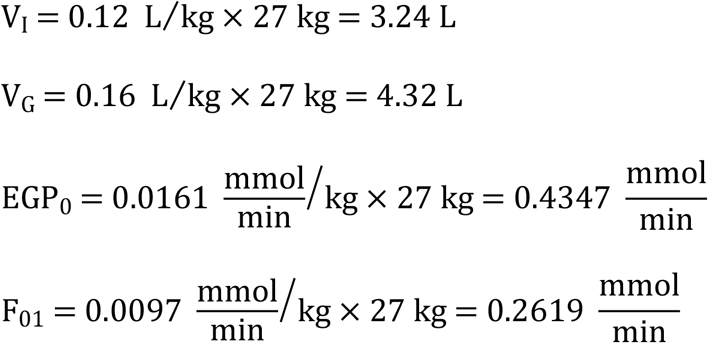

The same thing applied for patients 2 and 3 with body weight of 26.3 kg and 49.9 kg, respectively.

## Results and Discussion

### Meal Simulation in Glucose-Insulin Reaction

In-silico work was done to analyse the improved Hovorka equations with the presence of meal disturbances. The amounts of meals consumed can affect the BGL. Meals that contained carbohydrates (CHO) are broken down into glucose. The absence of sufficient insulin hormone in people with T1D increases BGL. This information will be beneficial in controlling BGL upon which the patients can inject insulin according to their body’s needs. If excess insulin is infused, patients may experience hypoglycaemia (low BGL) [12]. The calculations of the rate of meals can be referred to Daud et al. [10]. The total daily dose of insulin required can serve as a guide for the people with T1D to manage their BGL [9]. In-silico works with three meals were performed for each patient. All patients received bolus insulin 30-minute before every meal. Short acting insulin was infused earlier via the subcutaneous method due to the time lag for insulin to take effect [13]. Table 6 summarises the amount of bolus insulin administered for all patients in 24-hour. Bolus insulin administered was found using a trial and error to achieve optimum BGL possible.

### Simulation Results for Patient 1

Figure 2 shows the BGL profile against time for clinical and in-silico works in 24-hour for patient 1. The simulation starts at 05:00. During the fasting stage (no meal disturbances) in clinical work, patient 1 recorded BGL value of 15.8 mmol/L, whereas in-silico work was 6.9 mmol/L. Although the initial values of BGL for both clinical and in-silico works are different, there is a similarity in BGL trends during breakfast, lunch, and dinner time. As seen in Figure 2, patient 1 in clinical work experiences hyperglycaemia most of the day and only achieves normoglycaemia during breakfast time. In contrast, the BGL for patient 1 in in-silico work is able to achieve normoglycaemia 73% of the time.

**Figure 2.**
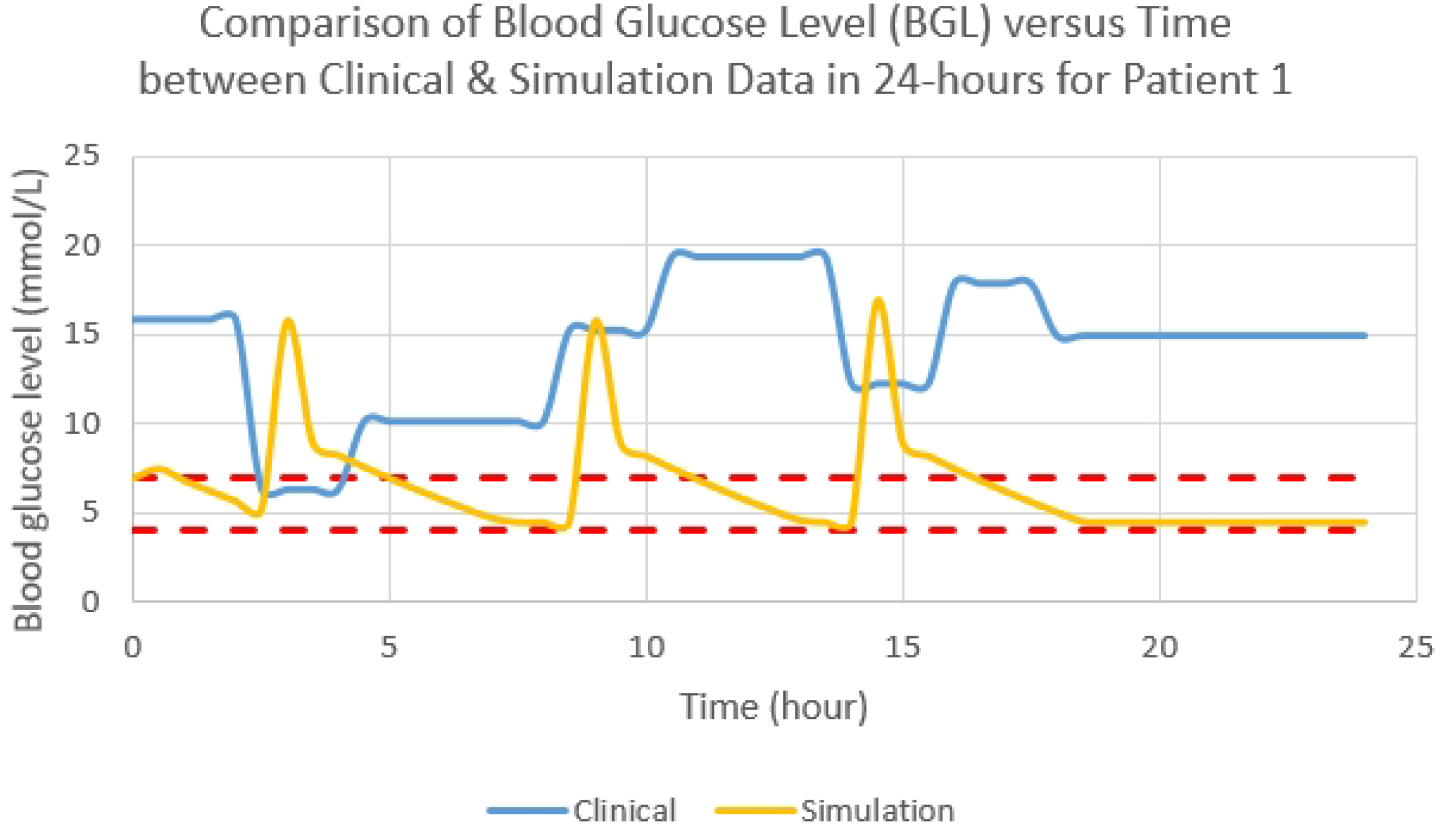
BGL profile in 24-hour for patient 1

### Simulation Results for Patient 2

Simulation in the presence of meal disturbances for patient 2 with a body weight of 26.3 kg was then executed as shown in Figure 3. It shows the BGL profile against time for clinical and in-silico works in 24-hour in which the BGLs recorded at the fasting stage are 13.2 mmol/L and 6.9 mmol/L, respectively. Based on Figure 3 and Table 4, patient 2 requires less bolus insulin to control BGL than the other two patients since the BGL deviates slightly above the normoglycemic range throughout the day but is more stable. Moreover, patient 2 in the in-silico work manages to stay in the normoglycemic range for 85% of the time compared to clinical work, which is merely 31%.

**Figure 3.**
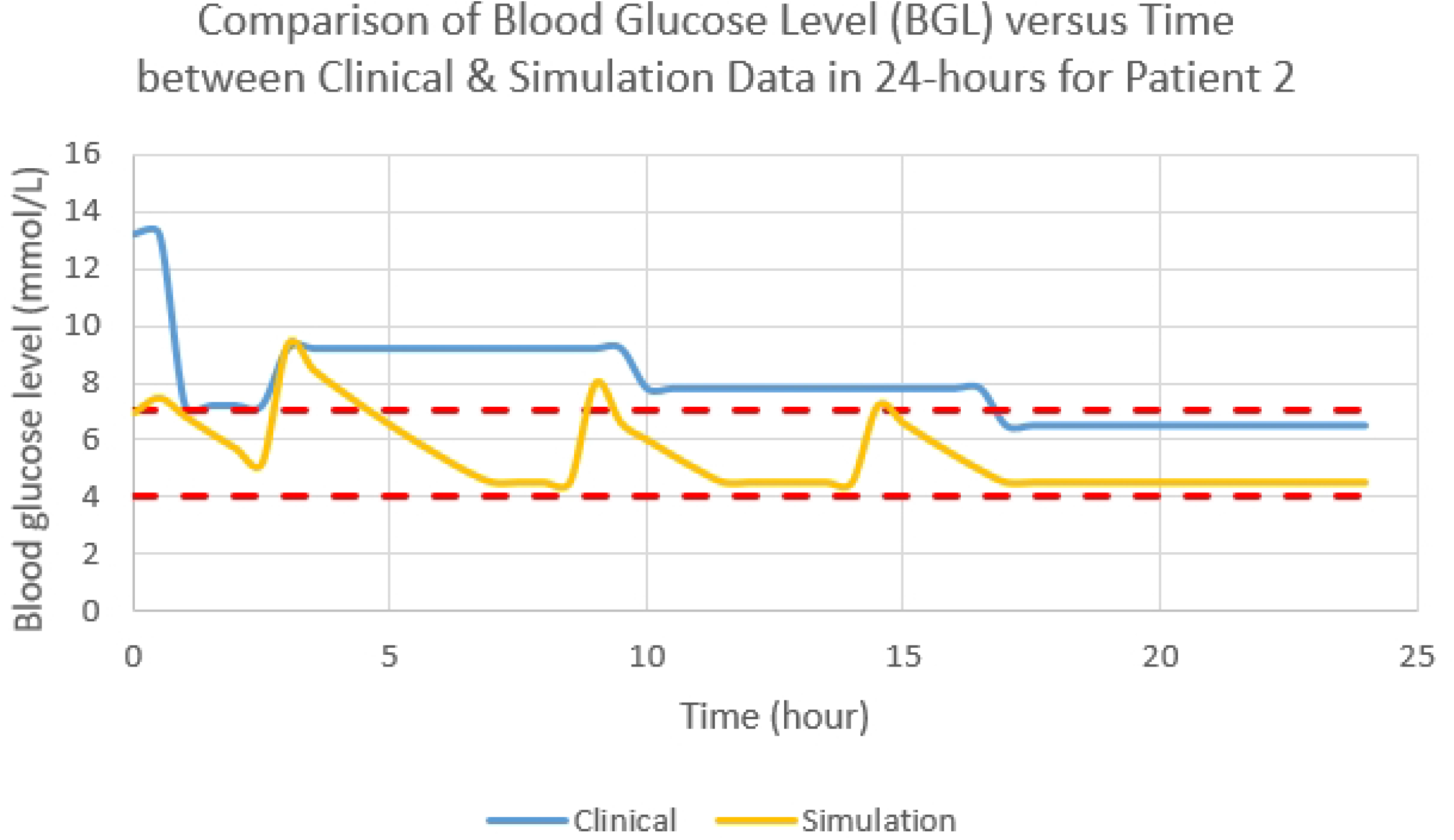
BGL profile in 24-hour for patient 2

### Simulation Results for Patient 3

The third simulation involved patient 3 with a body weight of 49.9 kg, which was done in the same manner as the previous two patients. Figure 4 shows the BGL profile against time for clinical and in-silico works in 24-hour for patient 3. Initially, BGLs recorded for clinical and in-silico works are 15.7 mmol/L and 5.0 mmol/L, respectively. Throughout the day, the patient’s BGL in the in-silico work experiences a similar trend with the clinical work and barely achieves normoglycemic range. For Patient 3, the time duration spent in the normoglycemic range was only 16% as in the in-silico work. Although the BGL in the in-silico work has better performance than in the clinical work, improvement should be done to increase more time duration spent in the normoglycemic range.

**Figure 4.**
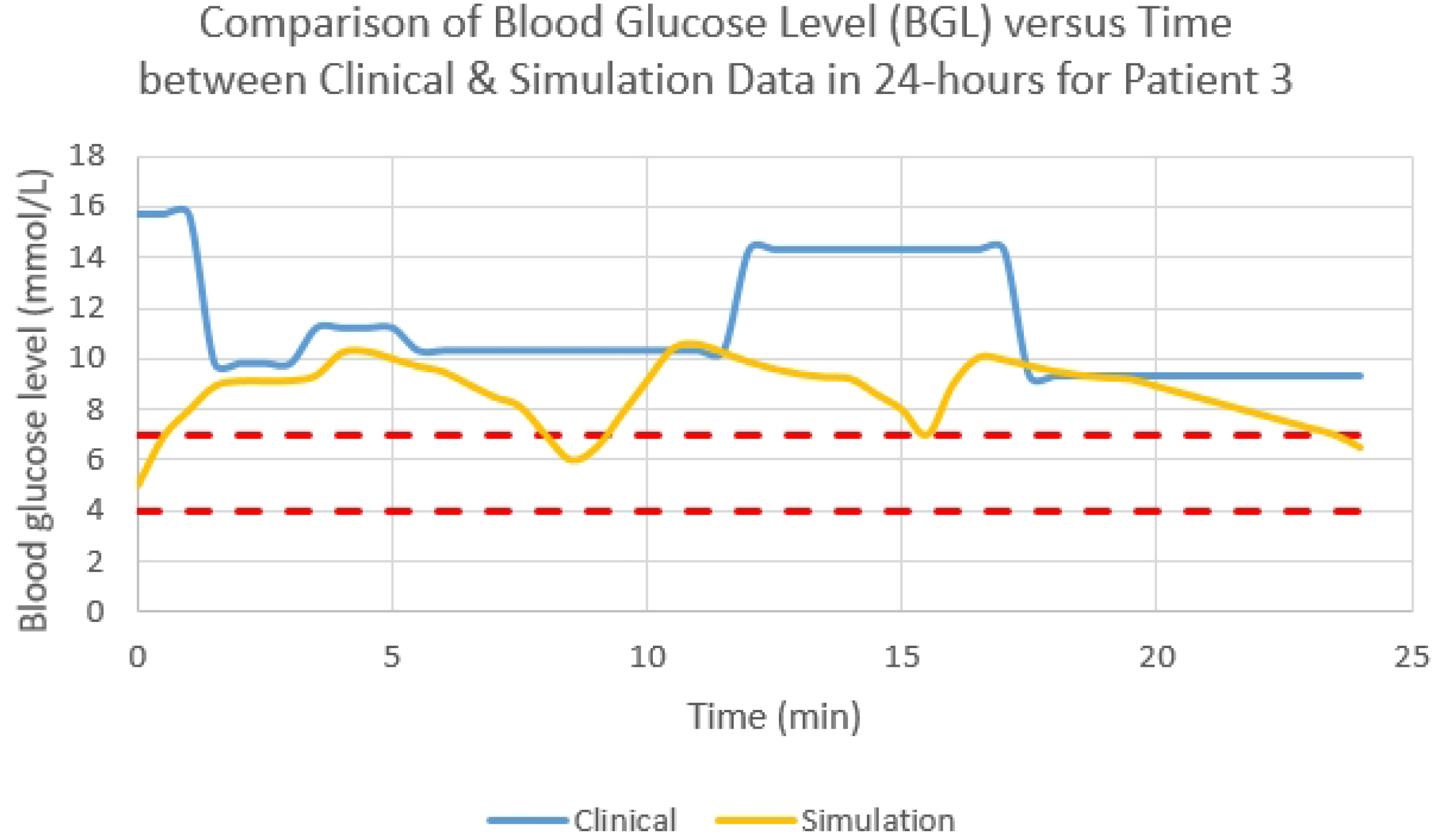
BGL profile in 24-hour for patient 3

Overall, all patients in the clinical works experienced hyperglycaemia most of the time in a day based on the BGL profile in 24-h. They also barely achieved the normoglycemic range during both pre-meal and post-meal. On the contrary, all patients showed improvement in BGL for the in-silico works. Large amount of exogenous insulin was infused during breakfast time compared to the rest of the day as insulin sensitivity was higher in the morning [14]. Table 7 shows regression statistics between clinical and in-silico works for all three patients. Both the clinical and in-silico works have p-values less than 0.05, thus indicate the simulation work using the improved Hovorka equations is acceptable to predict the BGL for people with T1D.

Since patients only recorded BGL a few times in a day, i.e., every pre-meal and post-meal intakes in clinical works, there is a possibility for their BGL to be in the normoglycemic range at other times. Moreover, high BGL in clinical works may be due to stress and physical activities. However, these factors are currently neglected in the improved Hovorka model equations. In future works, for better comparison of BGL between clinical and in-silico works, patients should be equipped with continuous glucose monitoring (CGM) device to track BGL continuously. Further works should also include a comparison between simulation results from Hovorka model equations and improved Hovorka model equations for model verification purposes. Perhaps, the current improved Hovorka equations can also be compared with other established process simulator such as UVA-Padova simulator.

## Conclusions

In this study, the improved Hovorka equations were presented based on Hovorka model (4) to study the effect of insulin input and meal disturbances on blood glucose level for people with T1D. The results showed an increased time spent in the normoglycemic range for all three patients in the in-silico work using the improved Hovorka equations compared to the clinical work. Thus, the improved Hovorka equations are proven reliable in simulating BGL with meal disturbances.

## Acknowledgements

The authors wish to acknowledge and extend their gratitude to the Ministry of Higher Education, Malaysia, and Research Management Centre, UiTM for the financial support given under Grant No: 600-RMC/GPK 5/3 (222/2020). The authors would also like to thank the School of Chemical Engineering, College of Engineering, UiTM and the Paediatric Department, Faculty of Medicine, UiTM for all the support and assistance rendered. The authors sincerely acknowledge the participation of people with T1D in this research project.

## Authorship Disclosure Statement

Nur Amanina Mohd Sohadi conducted the programming works. Ayub Md Som, Noor Shafina Mohd Nor, Sherif Abdulbari Ali and Mohd Aizad Ahmad analysed the data and supervised the overall research works. Nur Amanina Mohd Sohadi and Ayub Md Som wrote the paper. All authors have no conflict of interest.

